# Artificial light is the main driver of nocturnal feeding by the Rock dove (*Columba livia*) in urban areas

**DOI:** 10.1101/621508

**Authors:** Lucas M. Leveau

## Abstract

Artificial light at night (ALAN) is one of the most extreme alterations of urban areas, which drives nocturnal activity by diurnal species. Although the Rock Dove (*Columba livia*) is a common species in urban centers worldwide known to have nocturnal activity in urban areas, it is unknown what is the role of ALAN in its nocturnal activity. Moreover, studies that address the relationship between ALAN and nocturnal activity of diurnal birds are scarce in the Southern hemisphere. The objectives of this study were: 1) to evaluate the extent of nocturnal activity in the Rock Dove in large cities of Argentina; and 2) to analyze the influence of ALAN, pedestrian traffic and car traffic on the nocturnal activity in two cities, Buenos Aires and Salta. I visited the most urbanized areas of five large cities in Argentina, and surveyed lighted streets once after 30 minutes after sunset. In Buenos Aires and Salta, I compared environmental conditions between sites were doves were seen feeding with random sites were doves were not recorded feeding. Nocturnal feeding of the Rock Dove was recorded in three of five cities surveyed. ALAN was positively related to nocturnal feeding activity in Salta and Buenos Aires. The results obtained suggest that urbanization promotes a nocturnal activity of the Rock Dove, which occurs in cities located in a vast range of altitudes and biogeographic contexts. Moreover, the nocturnal activity is mainly driven by ALAN, which probably alters the circadian rhythm of doves.

## Introduction

Urban areas are characterized by a temporal stabilization of environmental conditions, which is related with a temporal stabilization of species composition (Leveau 2018). For instance, artificial light at night has been postulated as the main driver allowing the colonization of night by diurnal species. Focusing on birds, which are the most studied animal taxa in urban areas (Magle et al. 2012; Nielsen et al. 2014), several studies showed an increased activity at night for diurnal species in areas with the highest artificial light (Miller 2006; MacGregor Fors et al. 2011; Byrkjedal et al. 2012; Dominoni et al. 2013a; Da Silva and Kempenaers 2017). Conversely, artificial light at night could have negative impacts on nocturnal bird species, limiting their presence in urban areas (Weaving et al. 2011). As a result, the increased presence of diurnal birds at night and the local extinction of nocturnal species may lead to a temporal homogenization of bird communities (Leveau 2018).

Although the number of studies dealing with nocturnal activity of diurnal birds as a consequence of artificial light has increased in the last decade, most of them were conducted in the Northern Hemisphere, where the Blackbird (*Turdus merula*) was the most studied species (Leveau 2018). However, the Rock Dove (*Columba livia*), which is a cosmopolitan urban dweller (Aronson et al. 2014; Leveau and Zuria 2017) and has been recorded feeding at night in urban areas of Europe (Luniak 2004), has been scarcely studied regarding its relationship with artificial light.

Diurnal bird activity at night has been recorded as singing, feeding or both (Dominoni et al. 2013a, Da Silva and Kempenaers 2017, MacGregor Fors et al. 2011). Whereas bird singing at night may alter their breeding cycle with possible consequences on bird population dynamics (Miller 2006), bird feeding at night may have more profound consequences on interspecific relationships because nocturnal feeding can result in avoidance of interference competition with other diurnal species or the depleting of resources for nocturnal species (MacGregor Fors et al. 2011). For example, nocturnal omnivorous Rock Doves can deplete resources otherwise available for nocturnal rodents.

Given the lack of studies concerning the nocturnal activity of birds in the Southern Hemisphere and about the Rock Dove, the objectives of this study were: 1) to determine the presence of Rock Dove nocturnal feeding in the most urbanized areas in five of the ten largest Argentinian cities; and 2) to analyze the relationship between nocturnal feeding and artificial light, human presence and car traffic in Buenos Aires and Salta city.

## Material and Methods

The presence of nocturnal feeding was investigated in five of the most populated cities in Argentina (Table 1). In Salta, Buenos Aires, Mar del Plata and Rosario surveys were made between July and September 2016, corresponding to the austral winter. In Cordoba surveys were made during November 2015, corresponding to the austral spring. In all cities except Buenos Aires, surveys were made during one night in the most urbanized part of the city, which encompassed the surroundings of the central urban park and a pedestrian street. In Buenos Aires, given its larger size than the other cities, surveys were made three nights on the main avenues and the pedestrian street. Surveys started between 33 to 80 minutes after local sunset and lasted between 36 to 120 minutes.

**Table 1.**
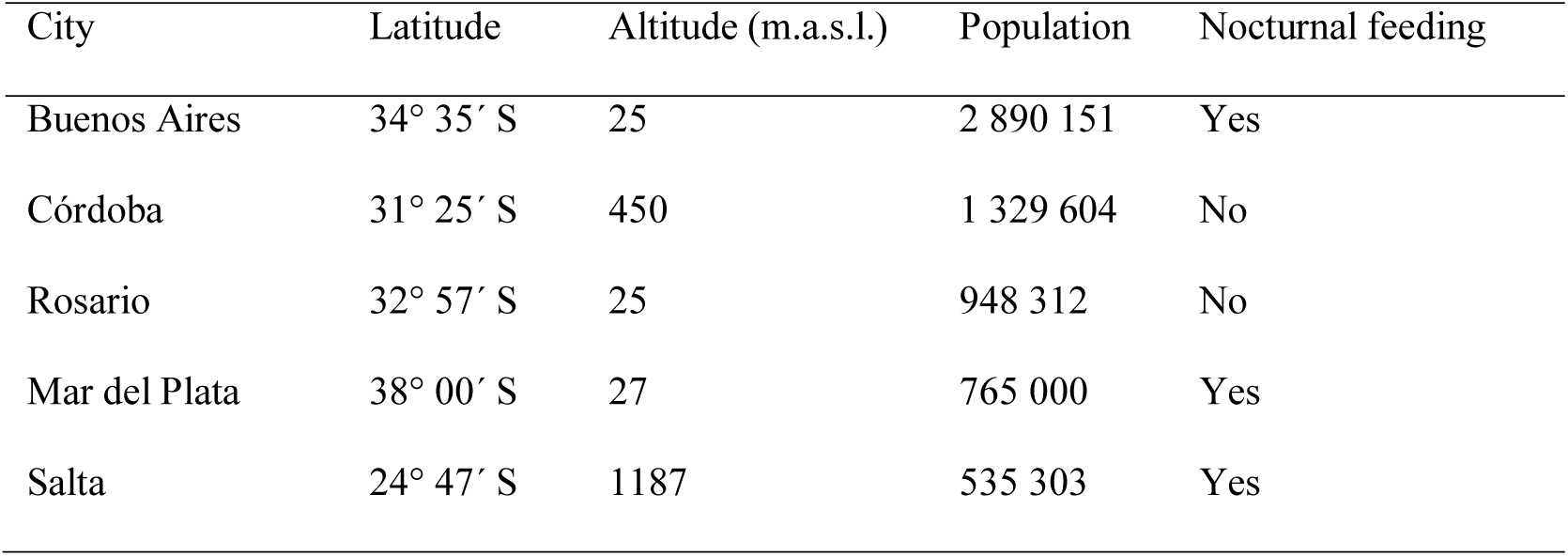
Characteristics of cities surveyed and presence of nocturnal feeding by Rock Doves in the most urbanized areas of five large cities of Argentina.

| City | Latitude | Altitude (m.a.s.l.) | Population | Nocturnal feeding |
| --- | --- | --- | --- | --- |
| Buenos Aires | 34° 35' S | 25 | 2 890 151 | Yes |
| Córdoba | 31° 25' S | 450 | 1 329 604 | No |
| Rosario | 32° 57' S | 25 | 948 312 | No |
| Mar del Plata | 38° 00' S | 27 | 765 000 | Yes |
| Salta | 24° 47' S | 1187 | 535 303 | Yes |

In cities with the most quantity of nocturnal feeding doves I located a similar number of random sites where doves were not seen feeding at night. In both sites light intensity, car and pedestrian traffic were measured once. Car and pedestrian traffic were calculated as the number of people and car passing during three minutes. In the case of pedestrian streets, there was no car traffic. Pedestrian traffic was considered an indicator of food for doves, because the greater the number of passing people, the greater the chance to obtain food for them intentionally or unintentionally. Car traffic was used as an indicator of noise, given that some studies suggested that noise is a determinant of diurnal bird activity at night (Fuller et al. 2007, Arroyo-Solis et al. 2013). Light intensity was measured with the cell phone application Lux Meter, using a cell phone Sony Xperia M. The cell phone was moved in all directions to obtain the mean value of lux, with an error of one lux.

### Statistical analysis

The relationship between dove nocturnal feeding and the environmental variables was analyzed by a Generalized linear mixed model (GLMM) with package lme4 and the function glmer in R (The R core team 2017). The response variable was the nocturnal feeding presence or absence, and binomial error structure was used. Given that records from two cities were used, city type was used as a random factor. Model selection was made from the full model with all environmental variables to the final model excluding non-significant variables and testing its significance with the anova function. Final model was compared against the null model, and plotted with the package visreg (Breheny and Burchett 2013). There was not a significant correlation between environmental variables (r < 0.70).

## Results

Doves feeding at night were observed in three of the five cities surveyed (Table 1). Buenos Aires and Salta had the highest number of nocturnal feeding events (eight and four, respectively), whereas in Mar del Plata only one nocturnal feeding event was recorded in the pedestrian street of downtown. In Buenos, during each feeding event between one to 16 doves were involved (mean = 7.5), whereas in Salta between one to three doves were seen. In Salta, nocturnal feeding was only observed in pedestrian streets, whereas in Buenos Aires was recorded in pedestrian streets and avenues.

There was a significant increase in the probability of nocturnal feeding with more light intensity (LRT = 8.27, P = 0.004; intercept = −1.84, slope = 0.04). A light intensity of more than 100 lux had the highest probability of nocturnal feeding occurrence (Figure 1).

**Figure 1.**
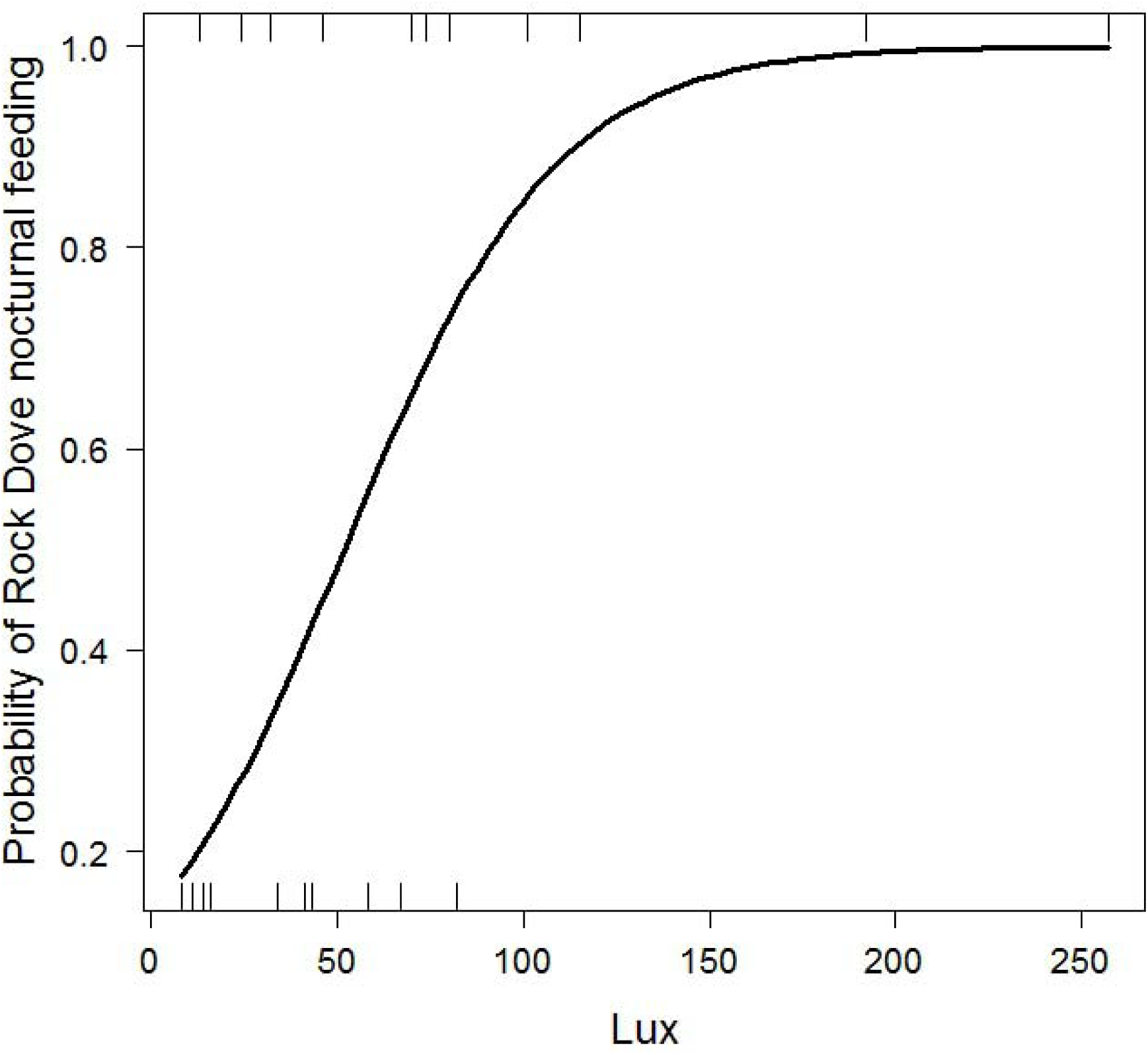
Probability of nocturnal feeding in the Rock Dove (*Columba livia*) in Salta and Buenos Aires in relation to light intensity (mean lux).

## Discussion

Results showed that nocturnal feeding of Rock doves occurred in more than half of cities surveyed, which had a notable biogeographic heterogeneity (from 24 masl to 1187 masl, and a difference of 14 degrees of latitude) suggesting that could be a generalized behavior among Latin American cities and worldwide judging by records reported by Luniak (2004) for Europe. However, the present study focused on large cities of more than 500 000 inhabitants, so the effect of urbanization on nocturnal dove behavior need to be studied in smaller cities. On the other hand, much more study is needed to elucidate why the nocturnal feeding of the Rock Dove is more likely in some large cities and absent in others. Possible explanations may be related to the levels of artificial light at night and the availability of discarded food by humans. Indeed, in three cases of nocturnal feeding in Buenos Aires two people were actively involved in feeding doves, whereas in one case doves were feeding on discarded food in a garbage container. Moreover, cities with elevated numbers of doves may encourage nocturnal feeding as a way to avoid intraspecific competition (Leveau 2018). Artificial light was the main predictor of nocturnal feeding by doves. This result agrees with other studies conducted in the northern hemisphere which focused on Passeriforms (MacGregor-Fors et al. 2011, Byrkjedal et al. 2012, Dominoni et al. 2013a, Stracey et al. 2014, Russ et al. 2014). However, our study showed that doves extended their activity after sunset according to Stracey et al. (2014) and Russ et al. (2014), whereas other studies only found a nocturnal activity just before twilight (Byrkjedal et al. 2012, Dominoni et al. 2013a). Regrettably, our survey did not cover the entire night, therefore I could not determine if doves extended their activity just before twilight. Moreover, other studies that included the same species such as the Blackbird and the European Robin (*Erithacus rubecula*), did not find a significant effect of light intensity and activity before twilight (Ockendon et al. 2009; Clewley et al. 2016). Disagreements among studies may be related to differences in light intensity. For instance, our analysis spanned a mean light intensity between 8 and 257 lux and was conducted in a highly urbanized area composed by high buildings and commercial areas. Nevertheless, Ockendon et al. (2009) and Clewley et al. (2014) conducted their study in a less urbanized area, composed by houses with gardens and probably with less light intensity.

Experimental studies showed that the increased activity induced by artificial light in birds is related to the reduced expression of melatonin (Dominoni et al. 2013b). Melatonin is secreted by the pineal gland and has a central role in the vertebrate circadian (Arendt 2000, Dominoni et al. 2013b). Studies in humans revealed that melatonin secretion was significantly suppressed at minimum light intensities that ranged between 1 lux to 393 lux (Brainard et al. 1982, McIntire et al. 1989, Aoki et al. 1998). Moreover, Russ et al. (2014) found a significant extension of night activity by Blackbirds in urban conditions with a mean light intensity of 0.44 lux. Results obtained in our study showed that a mean light intensity higher than 100 lux had the highest probability of nocturnal feeding behavior. This information is not only important to mitigate the temporal homogenization of bird communities but also to diminish the light pollution in urban areas which have negative impacts on human health (Haim and Zubidat 2015). Moreover, future studies should incorporate continuous measures of light intensities to compare among studies or species.

In this analysis, pedestrian traffic was used as an indicator of food availability for doves. However, this variable did not affect the nocturnal activity of doves. Other variables indicating food availability and not considered in this study, such as the number of garbage containers or restaurants may be more related to dove nocturnal feeding. On the other hand, noise was claimed by several studies as determinant of nocturnal singing in birds (Fuller et al. 2007, Arroyo-Solis et al. 2013). However, this study showed that nocturnal feeding by doves did not respond to car traffic, a proxy of noise.

## Conclusions

The results obtained showed that nocturnal activity by Rock Doves is common in Argentina, considering large cities. The nocturnal activity was recorded in cities located in a vast range of altitudes and latitudes. However, more studies are needed to know what environmental factors drive the biogeographical variation of nocturnal activity. For example, to analyze what city characteristics promote nocturnal bird feeding in the Rock Dove. At a local scale, our study showed that artificial light was the main driver of nocturnal feeding. Other factors such as food availability deserve more attention in future studies.

## Acknowledgements

I thank the help and support of Lucia Gonzalez Salinas.

